# Impact of Long-Term Cover Cropped Organic Farming Practices on the Development of Disease Suppressive Soils

**DOI:** 10.1101/2025.07.28.667314

**Authors:** Nahom Taddese Ghile, Sougata Bardhan, Frieda Eivazi, Chung-Ho Lin

**Affiliations:** Center for Agroforestry, Department of Forestry, University of Missouri, Columbia, MO, USA; Department of Forestry, School of Natural Resources, University of Missouri, Columbia, MO, USA; Department of Environmental Sciences, College of Agriculture Tennessee State University, Nashville, TN, USA; Department of Agriculture and Environmental Sciences, Lincoln University, Jefferson City, MO, USA

**Keywords:** Metabolomics, Cover Crops, Disease-Suppressive Soil, Rhizosphere, Organic Farming, UPLC-HRMS

## Abstract

Numerous field studies have reported the disease-suppressive properties of the system in organic farming and cover crop practices. This research examined how long-term organic farming with cover-cropped systems enhance soil disease suppressiveness and changes the chemical composition of the rhizosphere related to disease resistance. The study proposed that the interactions between plant exudates and helpful microorganisms in these systems create conditions that are less favorable for harmful pathogenic organisms, resulting in soil that suppresses disease. To achieve the goals, we used an untargeted metabolomics approach to compare chemical profiles from long-term certified organic fields, including different cover crop termination methods like flail and rotary mowing, to conventionally managed plots. We analyzed rhizosphere soil extracts using ultra-high-pressure liquid chromatography combined with high-resolution mass spectrometry (UPLC-HRMS). We processed data with both MS-DIAL and XCMS, which included peak detection and feature grouping. We annotated metabolites using the MS-DIAL Public VS17 libraries for features derived from MS-DIAL and the METLIN database for features derived from XCMS. Our analysis identified over 2500 distinct soil compounds, showing that long-term cover cropping significantly changes the soil’s chemical environment. The treatments with cover crops showed a higher abundance and diversity of bioactive chemicals compared to the controls, and this enriched metabolic profile supported the formation of disease-suppressive soil.

## 1. Introduction

Organic farming systems are receiving greater attention for their ability to enhance environmental sustainability, soil fertility, and high-quality crops. Farmers are searching for effective and eco-friendly management techniques as consumer demand for goods produced organically increases. One of organic farming’s main principles is reducing the use of synthetic inputs by implementing techniques that strengthen the system’s natural resilience, particularly in the areas of soil fertility, disease prevention, and pest control. The use of cover crops is one such technique that stands out as an essential strategy that offers organic farming systems a number of benefits (Bashir et al., 2025). Numerous field studies have reported the disease-suppressive properties of the system in organic farming and cover crop practices (Schlatter et al., 2017). This focus on soil resilience comes from the basic idea of the soil-plant-human health connection. This idea states that the health of the soil is closely linked to the health of plants, which then affects the health of the animals and people who consume them (Wall et al., 2015). Healthy, living soil produces more nutritious crops and can also lower the spread of human pathogens. They provide ecosystem services that are vital for public health (Pepper, 2013). So, practices that improve soil health are not just agricultural goals; they are crucial for enhancing overall human well-being. This makes the principles of organic farming a key part of a more unified “One Health” approach.

Cover crops, which are cultivated mostly for their soil advantages and not yield, are significant contributors to building up soil health in organic farming systems. By adding plant biomass, cover crops increase soil organic matter (SOM), which improves the soil’s structure, water-holding capacity, and nutrient cycling (Song et al., 2024). Cover crop decomposition replenishes the soil with essential nutrients, which could significantly reduce the need for outside organic inputs. Further, cover crops can suppress weeds, break disease and pest cycles, and hold soil in place, all of which add to farm resilience in general (Koudahe et al., 2022). The pH and nutrient content of the soil can also be impacted by the choice of particular cover crops and how they are terminated. For example, because legume cover crops release basic cations during decomposition, they can raise the pH of the soil. Flail mow and rotary mow termination can also have varying impacts on yields of subsequent crops and weed suppression. It is necessary to promote inherent disease resistance in the soil ecosystem due to the extensive limitations imposed on synthetic pesticides in organic management (Bashir et al., 2025). The development of disease-suppressive soils is one of the essential components of this inherent resilience. Disease-suppressive soils are those where, despite the presence of the pathogen and susceptible host, the microbial community and physicochemical characteristics of the soil reduce the incidence or severity of soilborne plant disease (Mendes et al., 2013). Soils that suppress disease are important in organic farming. The natural ability of soil to inhibit pathogens offers a sustainable and non-polluting method of disease control in situations where synthetic pesticides are not available or allowed. Suppression of disease can be separated into two general classes: general suppression, which is founded on total microbial biomass and activity and thus provides control over a broad range of pathogens, and specific suppression, which is characterized by the presence of high concentrations of individual microorganisms that are antagonistic to particular target pathogens (Scholberg et al., 2010). General suppression can be improved by SOM enhancement through cover cropping and organic input use, which stimulates beneficial microbial populations that compete with or antagonize pathogens. Specific suppression, while often requiring more specialized management, can also provide rigorous protection against specific diseases. Disease-suppressive soil formation is a multifactorial process that relies on soil amendments, cultural practices (cover crop and crop rotation), soil microbial flora, and plant-pathogen interactions (Song et al., 2020). Understanding and managing these factors is crucial for maintaining healthy and productive organic farming systems.

The factors behind this suppression are complicated and mainly involve chemical and biological interactions in the rhizosphere. These interactions include competition for nutrients, parasitism, and the release of various bioactive molecules. Beneficial microorganisms demonstrate sophisticated competitive strategies, including the production of high affinity siderophores that effectively sequester iron from the soil environment, thereby limiting pathogen access to this essential nutrient. Additionally, these microorganisms synthesize lytic enzymes such as chitinases, which target and degrade pathogen cell wall components (Berendsen et al., 2012). A major mechanism is antibiosis, where soil microbes release specific antifungal and antibacterial compounds. Notable examples are the production of 2,4-diacetylphloroglucinol (DAPG) by *Pseudomonas fluorescens* and other broad-spectrum antibiotics such as phenazines, polyketides, and lipopeptides (Mendes et al., 2013; Raaijmakers and Weller, 1998). Additionally, some beneficial bacteria block pathogen virulence through “quorum quenching”. They do this by breaking down N-acyl-homoserine lactones (AHLs), which are the signaling molecules pathogens use to organize their attack (Grandclément et al., 2016). Plants also play a direct role by releasing bioactive root exudates like flavonoids and strigolactones. These substances can selectively attract protective microbes to the rhizosphere (Hassan and Mathesius, 2012). This complex chemical communication network, mediated by numerous small molecules, forms the biochemical foundation of soil disease suppression systems. Therefore, understanding this molecular landscape is crucial for managing soil health and provides a strong reason for the following chemical profiling and metabolomic analysis.

Advanced analytical tools are needed to elucidate the mechanisms behind the complex interactions between cover cropping practices and the formation of soil disease-suppressive soil. Metabolomics, the global analysis of small molecular-weight compounds (metabolites) present in a biological system, has become an effective approach to characterizing the biochemical state of soils and understanding their functional attributes (Song et al., 2024). Metabolomics can provide insights about the active biochemistry reactions occurring in the soil ecosystem by measuring and detecting the diverse range of metabolites present in soil, including sugars, amino acids, organic acids, lipids, and secondary metabolites (Song et al., 2024).

Metabolomics has proven useful in defining soil quality, evaluating microbial activity, and characterizing responses to environmental change in soil health research (Withers et al., 2020). For example, Hayden et al. (2019) used soil metabolomics-proton nuclear magnetic resonance spectroscopy (^1^H NMR) and liquid chromatography–mass spectrometry (LC-MS) to distinguish between soils with different disease-suppressive capacities against *Rhizoctonia solani* AG8 in cereal crops, identifying macrocarpal as a putative biomarker of disease suppression. Metabolomics has also been applied to examine how agricultural management practices influence soil biochemical composition and microbial activity. For instance, studies have used metabolomics to evaluate how various land uses, like organic farming, affect soil metabolomes, and cover crops change the metabolite profiles of the soil. Cheng et al. (2018) presented evidence that the incorporation of biochar and the reduction of nitrogen fertilization in maize farming affected rhizosphere metabolomics. The importance of untargeted metabolomics for assessing microbial function and soil quality across a range of soil types and land uses was highlighted by Withers et al. (2020). These investigations demonstrate that the potential of metabolomics to offer meaningful insights into the intricate metabolic networks functioning in the soil and how they are associated with soil quality and disease suppression. This is especially true when high-resolution techniques like liquid chromatography-mass spectrometry combined with tandem mass spectrometry (LC-MS/MS) are used to identify a large number of known and unknown metabolites (Hayden et al., 2019).

While the role of organic practices in building soil health is well-established, the specific biochemical shifts that support the development of disease suppression over time remain poorly understood. A critical knowledge gap exists in connecting the duration and type of management to the soil’s functional chemical profile. This study, therefore, aims to elucidate these connections by using an untargeted metabolomics approach on soils from a long-term organic chronosequence. The primary objectives are: (1) to characterize the comprehensive soil metabolome to identify how it is shaped by the duration of cover crop management, and (2) to identify specific classes of metabolites, including fatty acids and bioactive secondary compounds, that are associated with mature organic systems. By linking these distinct chemical signatures to the mechanisms of general and specific suppression, this research seeks to uncover potential biomarkers of soil health and provide a mechanistic basis for how long-term cover cropping cultivates a disease-suppressive soil.

## 2. Materials and Methods

### 2.1. Site Description and Sample Collection

To compare soil metabolomes across different agricultural management methods, we collected sixty rhizosphere soil samples in sextuplicate from two main locations. One was Lincoln University’s Busby Farm, certified organic since 2013. The other was the conventionally managed Sanborn Field at the University of Missouri. At the Busby Farm organic site, we set up plots to test four fall cover crop termination methods: flail mowing, rotary mowing,bare tilling (with cover crop), and a bare tilled control (no cover crop). We assessed these treatments on plots in their first and fifth consecutive years of cover cropping to see how they changed over time. The cover crop mixture included hairy vetch (*Vicia villosa*), cereal rye (*Secale cereale*), and tillage radish (*Raphanus sativus*). For conventional reference, we obtained twelve samples from Sanborn Field, with equal samples taken from a rotational field (corn and wheat) and a monoculture corn plot. The complete experimental design is detailed in Figure 1.

**Figure 1.**
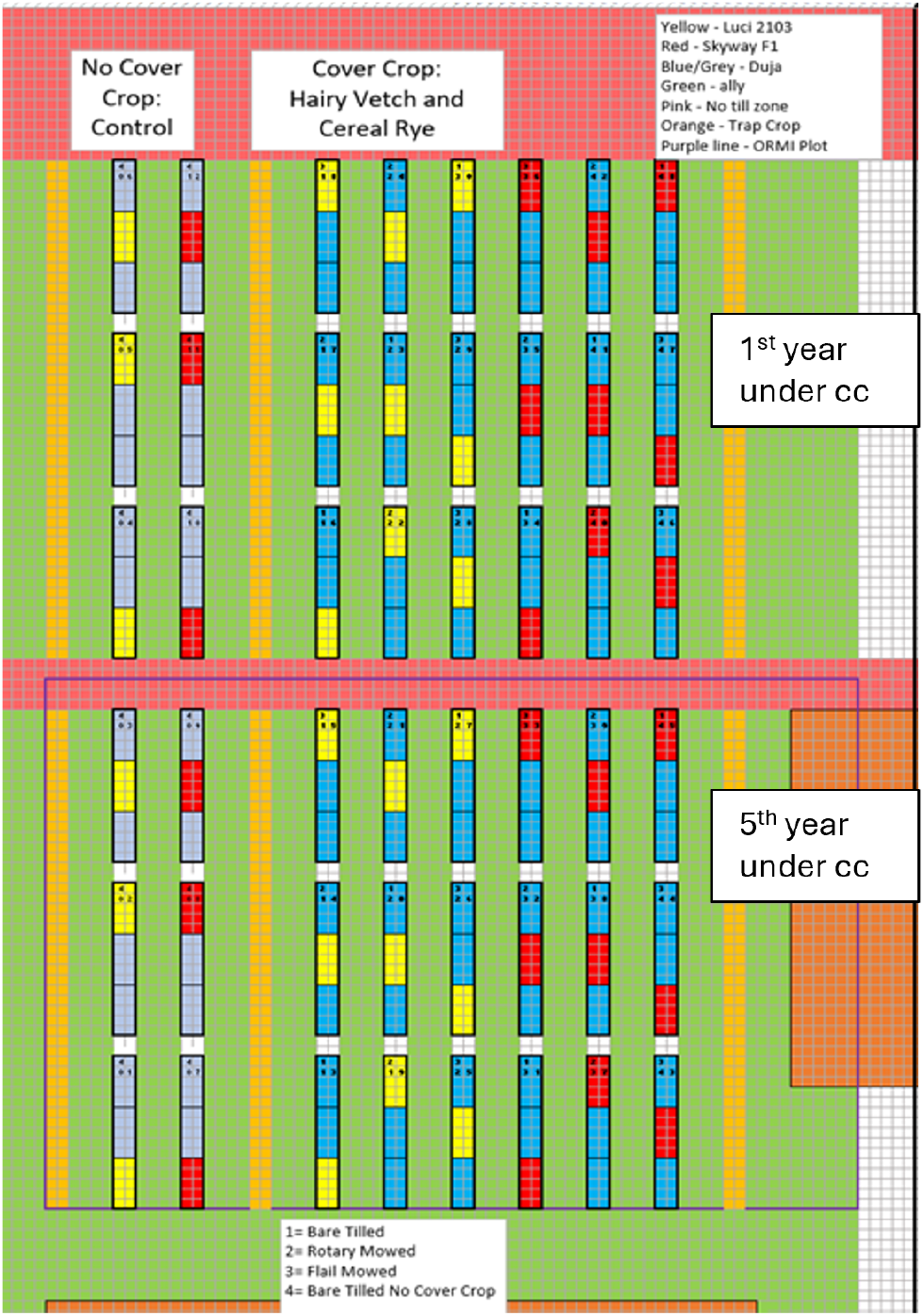
Experimental design at Lincoln University’s Busby Farm.

Immediately following collection, soil moisture was determined gravimetrically by oven-drying a subsample at 90°C for 24 hours. The remaining soil samples was stored at –20°C until analysis.

### 2.2. Metabolite Extraction

Soil metabolites were extracted from 2 g of dry weight equivalent soil using sequential methanol extractions. Each sample was transferred to a 15 mL centrifuge tube and extracted twice with 10 mL of analytical grade methanol. To prevent metabolite degradation, the soil methanol mixture was sonicated in an ice bath for one hour. After sonication, samples were centrifuged at 3,500 rpm for 10 minutes. After the first extraction, the supernatant was transferred to a 50 mL collection tube. A fresh 10 mL portion of methanol was added to the soil pellet for the second extraction cycle, which was performed under the same conditions. The combined supernatants, approximately 20 mL, were clarified by a final centrifugation step at 3,500 rpm for 10 minutes. Then, an 8 mL portion of the clear extract was transferred to a 15 mL glass tube, and the solvent was evaporated to dryness under a gentle stream of nitrogen. The dried residue was reconstituted in 1 mL of methanol and filtered through a 0.2 µm PTFE syringe filters before instrumental analysis.

### 2.3. UPLC-HRMS Metabolomic Profiling and Data Analysis

#### 2.3.1. Instrumental Analysis

Untargeted metabolomic profiling was performed using a Bruker maXis impact II quadrupole time-of-flight (Q-ToF) mass spectrometer coupled to a Waters Acquity Ultra Performance Liquid Chromatography (UPLC) system. Chromatographic separation was achieved on a Waters BEH C18 column (2.1 × 100 mm, 1.7 µm) maintained at 60 °C. Mobile phase A consisted of 0.1% formic acid in water, and mobile phase B was acetonitrile, delivered at 0.56 mL/min. The linear gradient program was as follows: starting at 5% B and ramping to 70% B over 30 minutes; increasing to 95% B from 30 to 33 minutes and holding at 95% B for 3 minutes; then returning to 5% B from 36 to 40 minutes to re-equilibrate the column. We conducted high-resolution mass spectrometry in both positive and negative electrospray ionization (ESI) modes. Key ESI source parameters included a nebulization gas pressure of 43.5 psi, a dry gas flow of 12 L/min, a dry temperature of 250°C, and a capillary voltage of 4000 V. Auto MS/MS spectra were collected for the top three precursors with a threshold of 10 counts, using mass-dependent collision energy. The instrument was auto calibrated with a post-run injection of sodium formate.

#### 2.3.2. Data Processing and Annotation

Data processing and annotation were carried out using two software platforms to analyze the metabolomic data. For initial untargeted profiling and statistical analysis, MS1 data were processed using the XCMS Online platform. Raw ion chromatograms underwent peak detection with the centWave algorithm. This was followed by feature grouping, retention-time alignment, and spectral extraction. The platform performed pairwise comparisons, flagging features with Welch’s t-test p-values below 0.01 and calculating fold changes. Tentative annotations were assigned by matching accurate ion masses with the METLIN database. To improve compound identification using fragmentation data, the raw data files were also processed with MS-DIAL (v4.9). The MS-DIAL workflow involved peak picking, deconvolution of MS2 spectra, and feature alignment. Data processing parameters included mass accuracy tolerances of 0.01 Da for MS1 and 0.05 Da for MS2, a minimum peak height of 1,000 amplitudes, and a retention time tolerance of 0.15 minutes for alignment. Putative annotations were made by matching the deconvoluted fragment spectra to public MS/MS libraries (MSMS-Public-Pos-VS17 and MSMS-Public-Neg-VS17) with an identification score threshold of 80%.

## 3. Results

### 3.1. Statistical Identification of Significant Metabolic Features

The untargeted UPLC-HRMS analysis of rhizosphere soil extracts successfully identified over 2,500 distinct metabolic features. To identify which of these features were significantly affected by the different agricultural management practices, we first employed a one-way analysis of variance (ANOVA). The results, visualized in a Manhattan plot, revealed hundreds of features with statistically significant differences (p < 0.05) across the treatment groups, with a subset showing highly significant p-values (Figure 2). To complement the ANOVA and control the false discovery rate (FDR), a Significance Analysis of Metabolites (SAM) was performed. This method identified 968 metabolic features as being significantly different among the treatments, with a low FDR of 4.6% at a delta value of 0.5 (Figure 3). Together, these statistical tests confirmed that long-term management practices induced substantial and significant changes in the soil metabolome, providing a robust basis for subsequent multivariate and targeted analyses.

**Figure 2.**
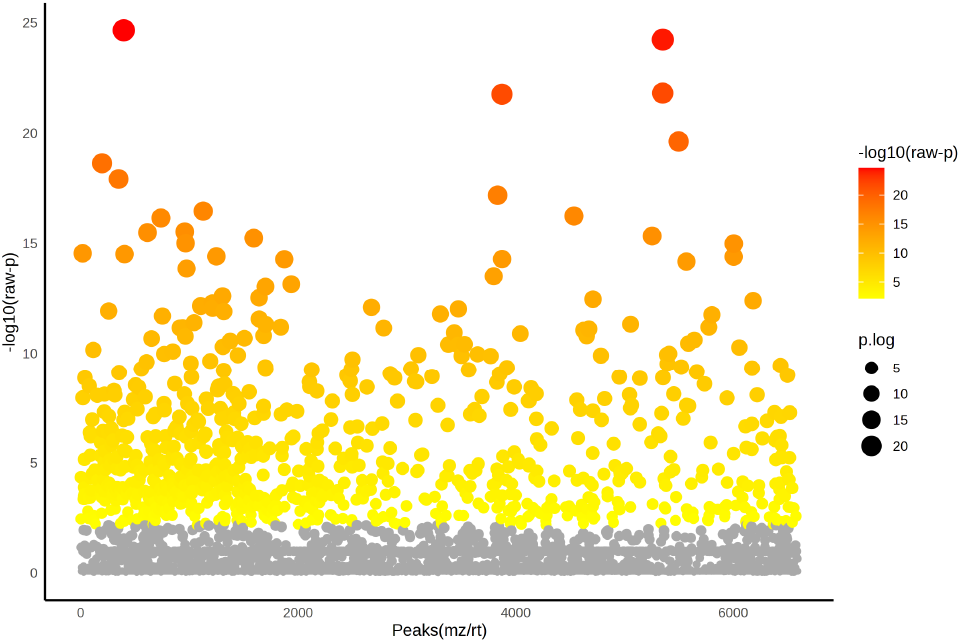
Manhattan plot of ANOVA results for all detected metabolic features across treatment groups. Each point represents a feature, with its vertical position indicating the level of statistical significance as −log10(raw p-value). Points colored yellow to red are considered statistically significant (p < 0.05), demonstrating that hundreds of metabolites were significantly altered by the different management practices.

**Figure 3.**
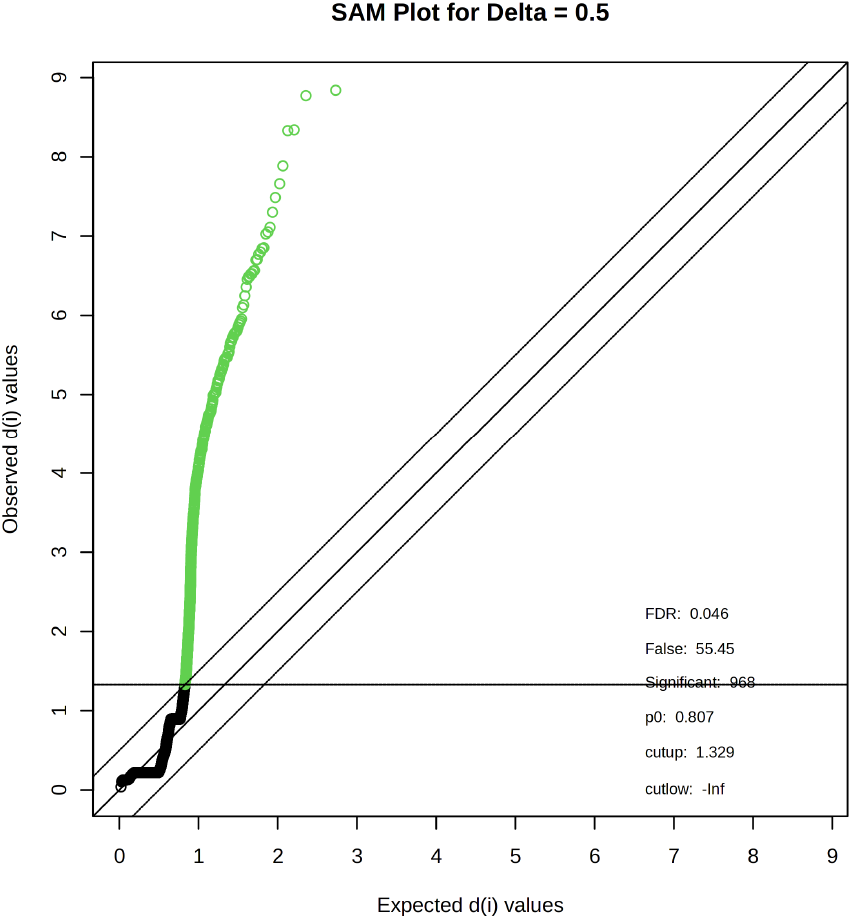
Significance Analysis of Metabolites (SAM) plot comparing observed vs. expected statistical scores. The green points represent the 968 metabolic features identified as statistically significant with a False Discovery Rate (FDR) of 4.6%. This analysis confirms that the number of significant features greatly exceeds what would be expected by chance.

### 3.2. Overall Metabolic Profile and Treatment Differentiation

To visualize the overall differences in metabolism driven by the most significant compounds, we performed a Partial Least Squares-Discriminant Analysis (PLS-DA) on the top 50 features identified by ANOVA. These features were annotated based on their accurate mass (MS^1^-level annotation). The scores plot shows a clear separation between the treatment groups (Figure 4). Notably, the mature cover-cropped organic systems (CC 5th year) cluster distinctly from the 1st year organic and conventional systems. This separation indicates that the length of covercrop use in organic management is a key factor in the metabolic profile. The heatmap of these same 50 metabolites further illustrates this pattern, showing a group of compounds that are highly abundant in the 5th-year cover crop treatment compared to all other groups (Figure 5).

**Figure 4.**
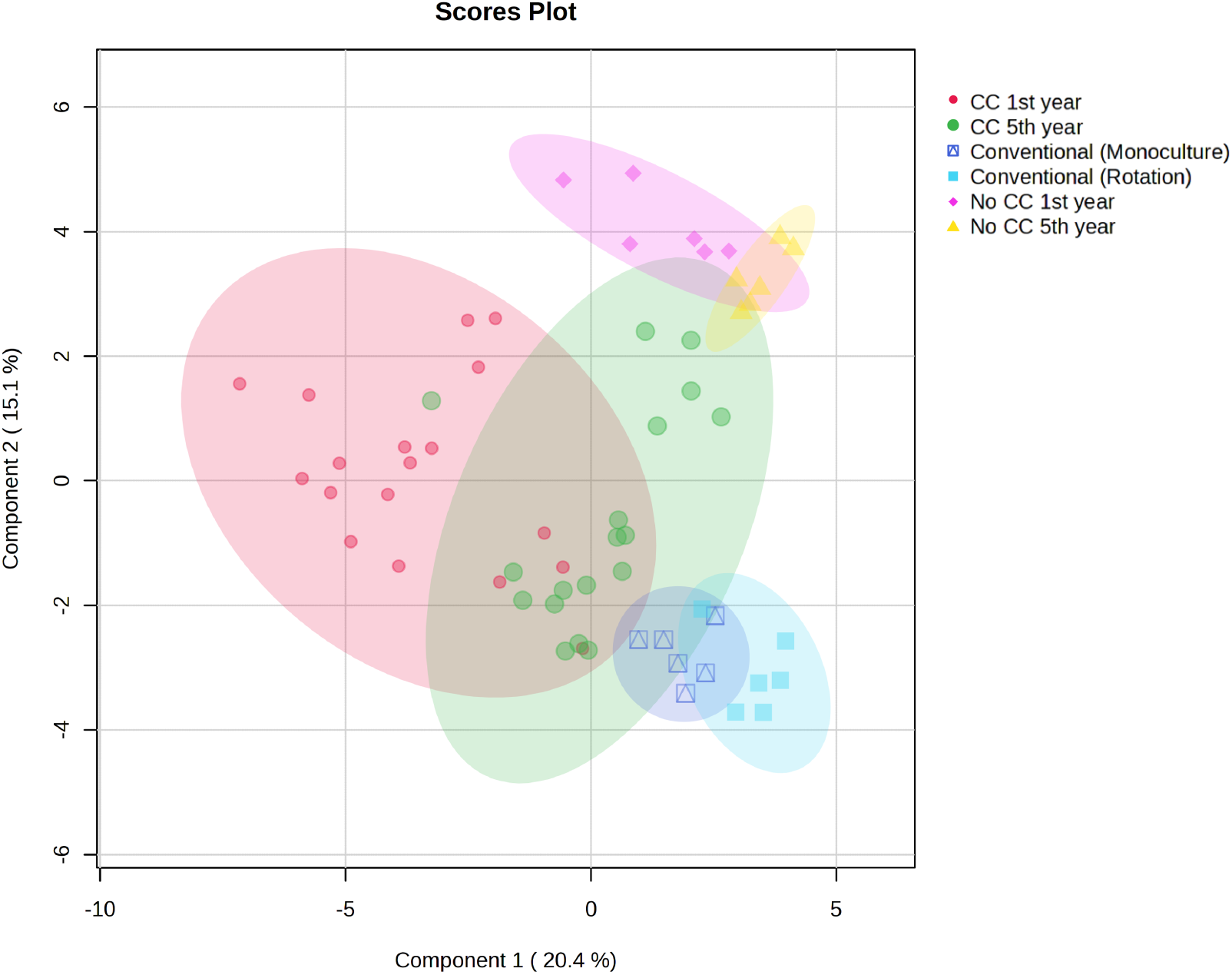
PLS-DA scores plot of the top 50 most significant metabolites (MS^1^ annotation). Each point represents a soil sample, and the ellipses represent the 95% confidence interval for each treatment group. The plot shows clear clustering by management practice and duration.

**Figure 5.**
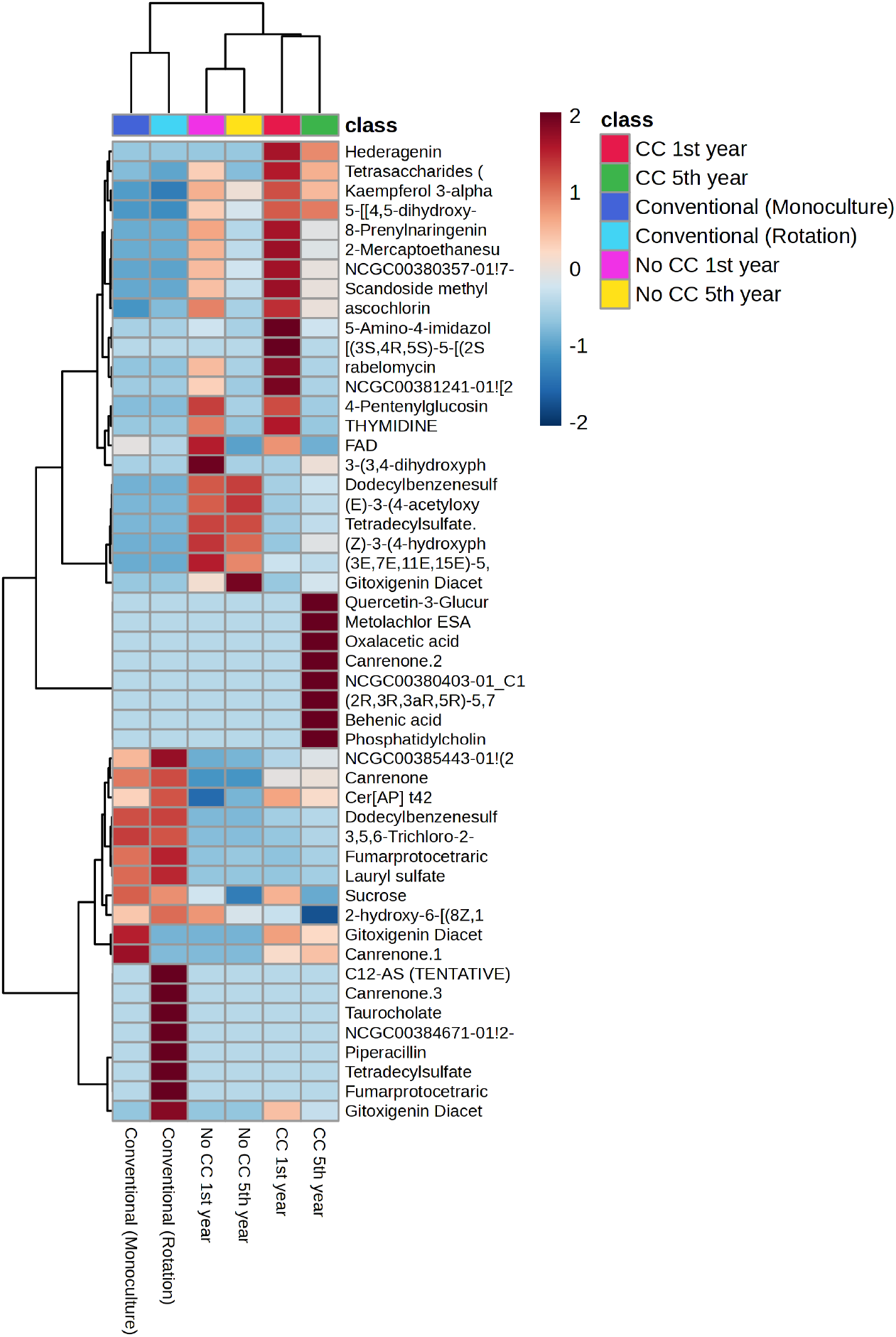
Heatmap illustrating the relative abundance of the top 50 most significant metabolites (MS^1^ annotation) across treatments. Each row represents a metabolite, and each column represents a treatment group. The color scale indicates normalized abundance, with red representing high abundance and blue representing low abundance. The analysis highlights a distinct block of highly abundant metabolites in the 5th-year cover crop (CC) treatment.

We also conducted a more targeted sPLS-DA on a curated list of known bioactive metabolites, which were annotated with higher confidence using fragmentation data (MS^2^-level annotation), with the exception of aconine, gossypin, picrotin, and sclareol (MS^1^). This analysis showed a clear and significant difference between the organic cover crop (CC) systems and the no-cover-crop (No CC) controls (Figure 6). While some overlap was noted between the 1st-year treatments, the 5th-year CC treatment formed a tight, distinct cluster. The differences became even clearer when we included conventional management systems in the analysis (Figure 7). In this context, the mature (5th year) organic CC system occupied a unique position in the plot, separating it from all other treatment groups.

**Figure 6.**
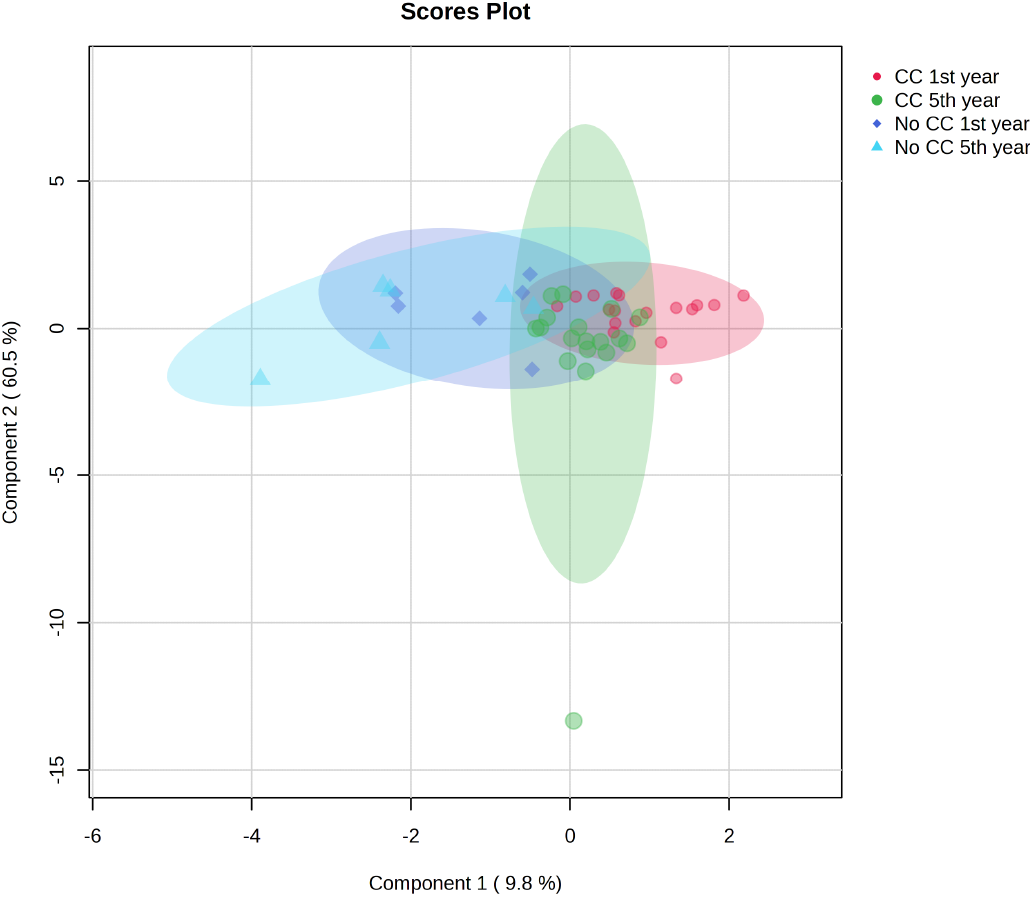
sPLS-DA scores plot of curated bioactive metabolites (primarily MS^2^ annotation) from organic systems.

**Figure 7.**
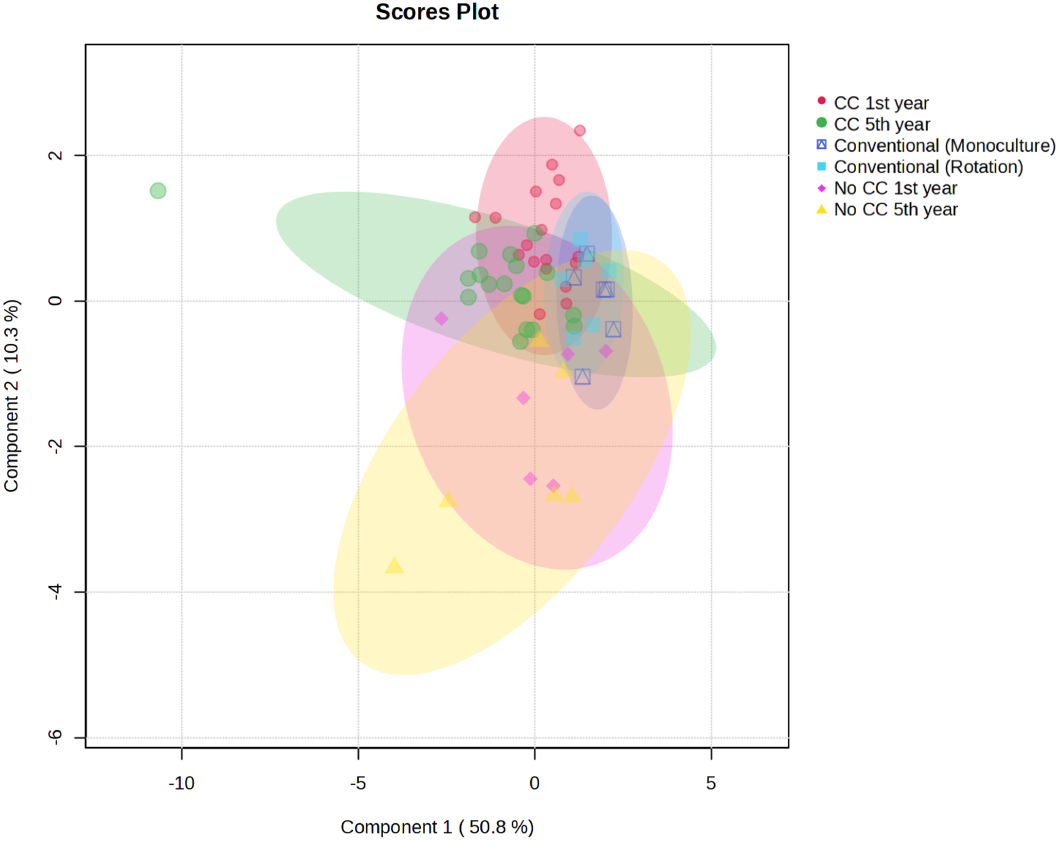
sPLS-DA scores plot of curated bioactive metabolites including conventional management systems.

### 3.3. Identification of Key Differentiating Metabolites

To rank the most influential compounds within the curated bioactive subset, a Random Forest classification model was used. This machine-learning approach ranked metabolic features based on their importance to classification accuracy (Mean Decrease Accuracy). This process successfully identified, from the curated key bioactive compounds, those most responsible for distinguishing the treatments. Among the top-ranking features were several fatty acids (pentadecanoic acid, eicosenoic acid) and the fatty acid amide oleamide (Figure 8), suggesting that major shifts in microbial community structure and signaling were primary drivers of the overall metabolic differences. A heatmap was subsequently generated to visualize the relative abundance of these key differentiating metabolites across all treatments (Figure 9). This provided a clear overview of the chemical shifts, revealing a distinct pattern of accumulation, particularly in the 5th-year CC treatment. A notable block of highly abundant compounds, visually represented by a strong red signature, distinguishes the mature cover crop system. This enrichment was especially apparent for known bioactive secondary metabolites and various fatty acids when compared to the relatively depleted profiles of the 1st-year CC, No CC, and conventional systems, highlighting a targeted chemical enrichment fostered by long-term cover cropping.

**Figure 8.**
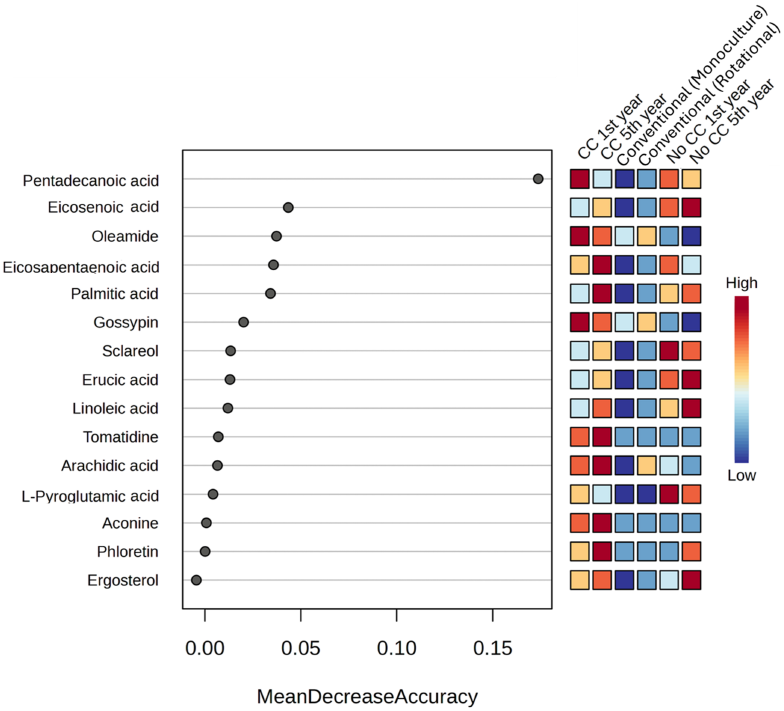
Random Forest analysis results. Features are ranked by their contribution to classification accuracy (Mean Decrease Accuracy). A higher value shows greater importance for the classification model. Pentadecanoic acid, eicosenoic acid, and oleamide were some of the most important features for distinguishing between management practices in this subset.

**Figure 9.**
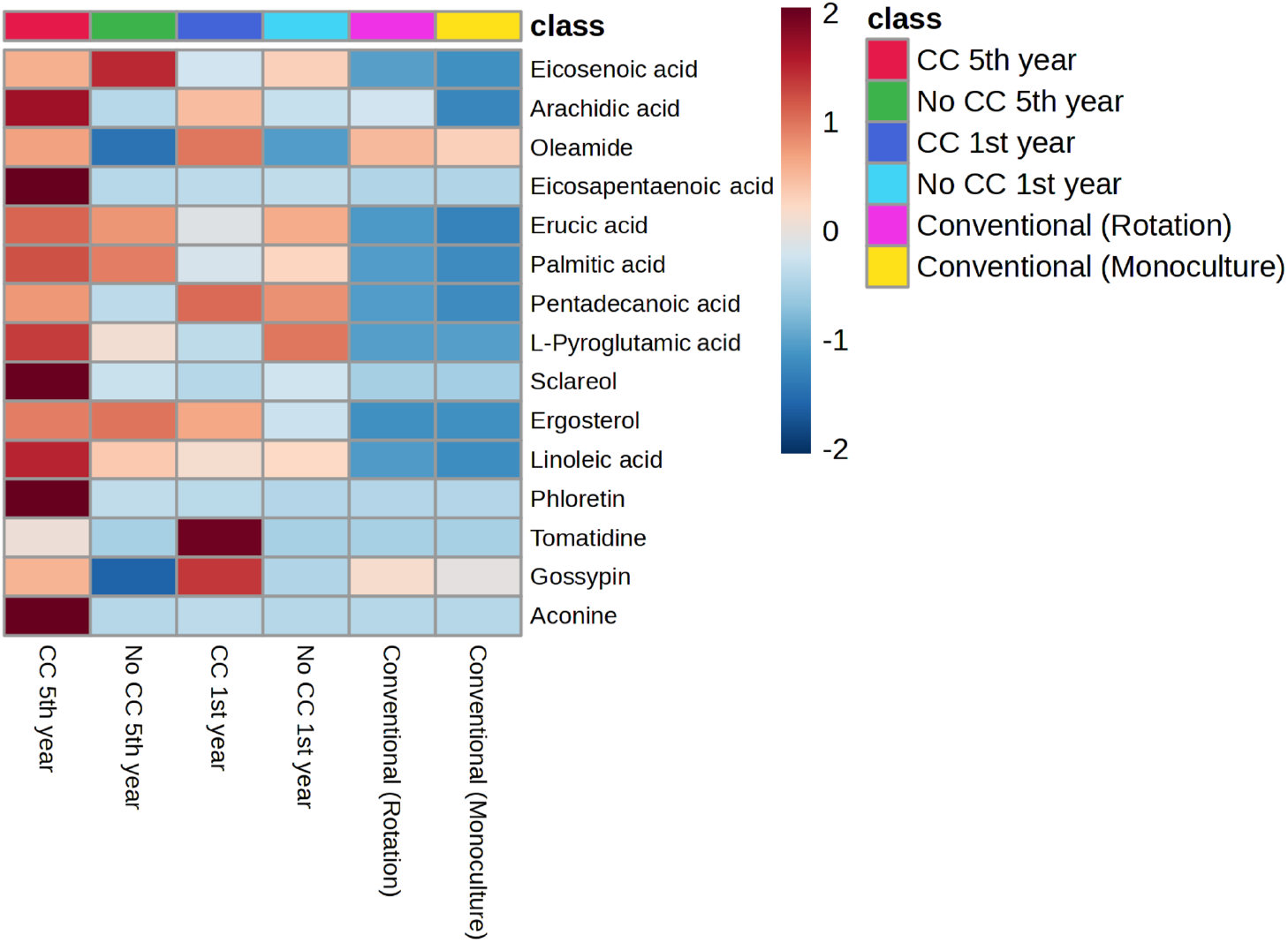
Heatmap illustrating the relative abundance of key selected metabolites (MS^2^ annotation, except for aconine, gossypin, picrotin, and sclareol) across treatments.

### 3.4. Dynamics of Specific Bioactive Compounds

From the statistically significant features, twenty compounds of known biological relevance were selected for detailed examination (Table 1). The majority of these compounds were identified with high confidence using fragmentation data (MS^2^), with the exceptions of aconine, gossypin, picrotin, and sclareol, which were annotated by accurate mass (MS^1^).

**Table 1:** Twenty compounds putatively identified in soil extract using metabolomics approach.

| Putatively identified compounds | RT (min) | Formula | Adducts | Theoretical mass (Da) | Observed mass (Da) | $\Delta m$ (mDa) |
| --- | --- | --- | --- | --- | --- | --- |
| Cordycepin | 18.3 | $C_{10}H_{13}N_5O_3$ | [M+H]<sup>+</sup> | 251.101839 | 251.107025 | 4.85 |
| Aconine | 21.7 | $C_{25}H_{41}NO_9$ | [M+H]<sup>+</sup> | 499.278132 | 499.271685 | 0.49 |
| Hesperidin | 23.3 | $C_{28}H_{34}O_{15}$ | [M-H]<sup>-</sup> | 610.189770 | 610.197645 | 1.34 |
| Gossypin | 32.02 | $C_{21}H_{20}O_{12}$ | [M+H]<sup>+</sup> | 480.090391 | 480.381215 | 0.97 |
| Picrotin | 0.979 | $C_{15}H_{18}O_6$ | [M+H]<sup>+</sup> | 310.105253 | 310.105095 | 0.09 |
| Fraxinol | 13.9 | $C_{11}H_{10}O_5$ | [M+H]<sup>+</sup> | 222.052823 | 222.056215 | 4.04 |
| Tomatidine | 10.12 | $C_{27}H_{45}NO_2$ | [M+H]<sup>+</sup> | 415.345030 | 415.344505 | 0.04 |
| Phloretin | 11.06 | $C_{15}H_{14}O_5$ | [M-H]<sup>-</sup> | 274.084124 | 274.088025 | 0.77 |
| Linoleic Acid | 31.28 | $C_{18}H_{32}O_2$ | [M-H]<sup>-</sup> | 280.240230 | 280.240725 | 0.09 |
| Ergosterol | 28.96 | $C_{28}H_{44}O$ | [M+H]<sup>+</sup> | 396.339216 | 396.339735 | 0.09 |
| Sclareol | 32.48 | $C_{20}H_{36}O_2$ | [M-H]<sup>-</sup> | 308.271530 | 308.271925 | 0.09 |
| L-Pyroglutamic Acid | 0.77 | $C_5H_7NO_3$ | [M-H]<sup>-</sup> | 129.042593 | 129.043325 | 4.51 |
| Pentadecanoic Acid | 33.63 | $C_{15}H_{30}O_2$ | [M-H]<sup>-</sup> | 242.224580 | 242.225325 | 0.09 |
| Palmitic Acid | 34.21 | $C_{16}H_{32}O_2$ | [M-H]<sup>-</sup> | 256.240230 | 256.240825 | 0.01 |
| Erucic acid | 33.81 | $C_{22}H_{42}O_2$ | [M-H]<sup>-</sup> | 338.318481 | 338.319105 | 1.22 |
| Palmitoleic Acid | 29.08 | $C_{16}H_{30}O_2$ | [M-H]<sup>-</sup> | 254.224580 | 254.225405 | 0.17 |
| Eicosapentaenoic Acid | 28.01 | $C_{20}H_{30}O_2$ | [M-H]<sup>-</sup> | 302.224580 | 302.224985 | 0.13 |
| Oleamide | 29.61 | $C_{18}H_{35}NO$ | [M+H]<sup>+</sup> | 281.271865 | 281.271685 | 0.49 |
| Arachidic Acid | 33.3 | $C_{20}H_{40}O_2$ | [M-H]<sup>-</sup> | 312.302831 | 312.303325 | 0.09 |
| Eicosenoic acid | 33.09 | $C_{20}H_{38}O_2$ | [M-H]<sup>-</sup> | 310.287181 | 310.288095 | 0.06 |

**Table 2:** Mechanisms of action for key bioactive compounds relevant to disease suppression.

| Compound | Chemical Class | Primary Mechanism(s) of Action | References |
| --- | --- | --- | --- |
| Hesperidin | Flavonoid | Signaling molecule; precursor to more active antimicrobials (hesperetin) via microbial biotransformation. | Choi et al. (2022) |
| Tomatidine | Steroidal Alkaloid | Detoxification product for specialized pathogens (*F. oxysporum*); retains antifungal activity against non-adapted fungi; can suppress host plant defenses. | Ito et al. (2004), Ito et al. (2007), Pareja-Jaime et al. (2008) |
| Sclareol | Diterpene | Direct antifungal (induces apoptosis, disrupts membranes); induces host plant defense responses; allelopathic effects. | Seo et al. (2012) |
| Phloretin | Dihydrochalcone | Potent allelochemical (involved in apple replant disease); direct antibacterial and antifungal activity. | Gallegos et al. (2016), Đorđić et al. (2021) |
| Ergosterol | Sterol | Biomarker for living fungal biomass, indicating an active and dynamic fungal community. | Montgomery et al. (2000) |

Analysis of these individual compounds showed several clear trends related to long-term cover cropping. A key finding was the significant accumulation of various known bioactive secondary metabolites in the cover-cropped plots over the five-year period. The amounts of the alkaloids aconine and tomatidine, the diterpene sclareol, and the dihydrochalcone phloretin increased significantly from the first year to the fifth year under cover cropping. In contrast, these compounds stayed near zero in the no-cover-crop controls. This demonstrates that their presence directly results from the cover crop management practice (Figure 10).

**Figure 10.**
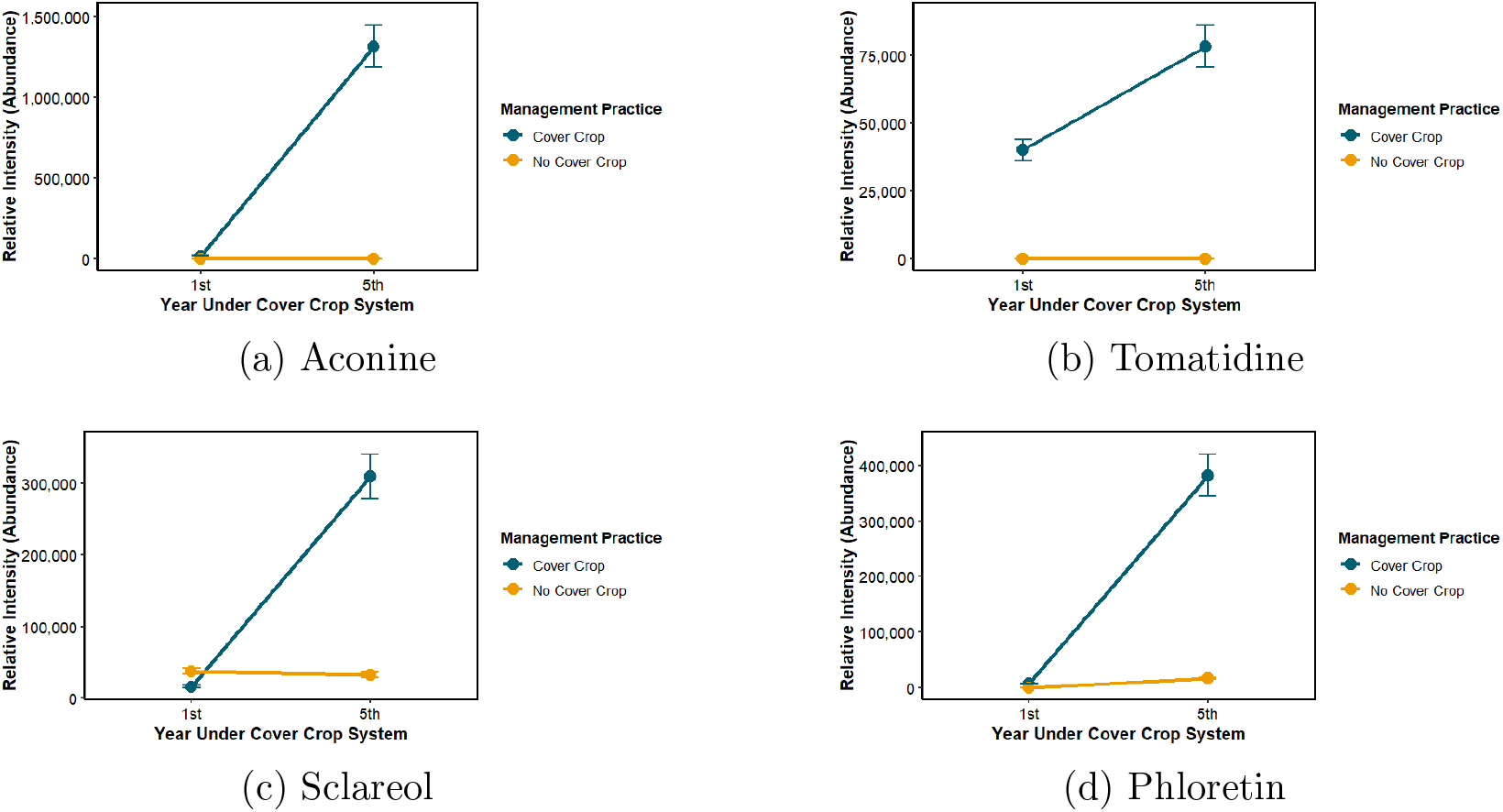
Relative intensity of select bioactive secondary metabolites. Aconine (A), Tomatidine (B), Sclareol (C), and Phloretin (D) all show significant accumulation from the 1st to the 5th year exclusively in the cover crop (CC) treatment.

Fatty acids are important markers of microbial biomass and community structure. They showed significant and varied changes. Long-chain fatty acids, like palmitic acid and arachidic acid, accumulated in the 5th-year CC treatment. This aligns with an increase in total microbial biomass due to a steady influx of organic matter. In contrast, fatty acids such as pentadecanoic acid and palmitoleic acid declined over time in both the CC and No CC treatments (Figure 11). This opposite trend hints at more complex changes in the microbial community. It may indicate a shift in the dominant microbial groups or a transition from fast-growing to slow-growing microbes as the soil ecosystem matured.

**Figure 11.**
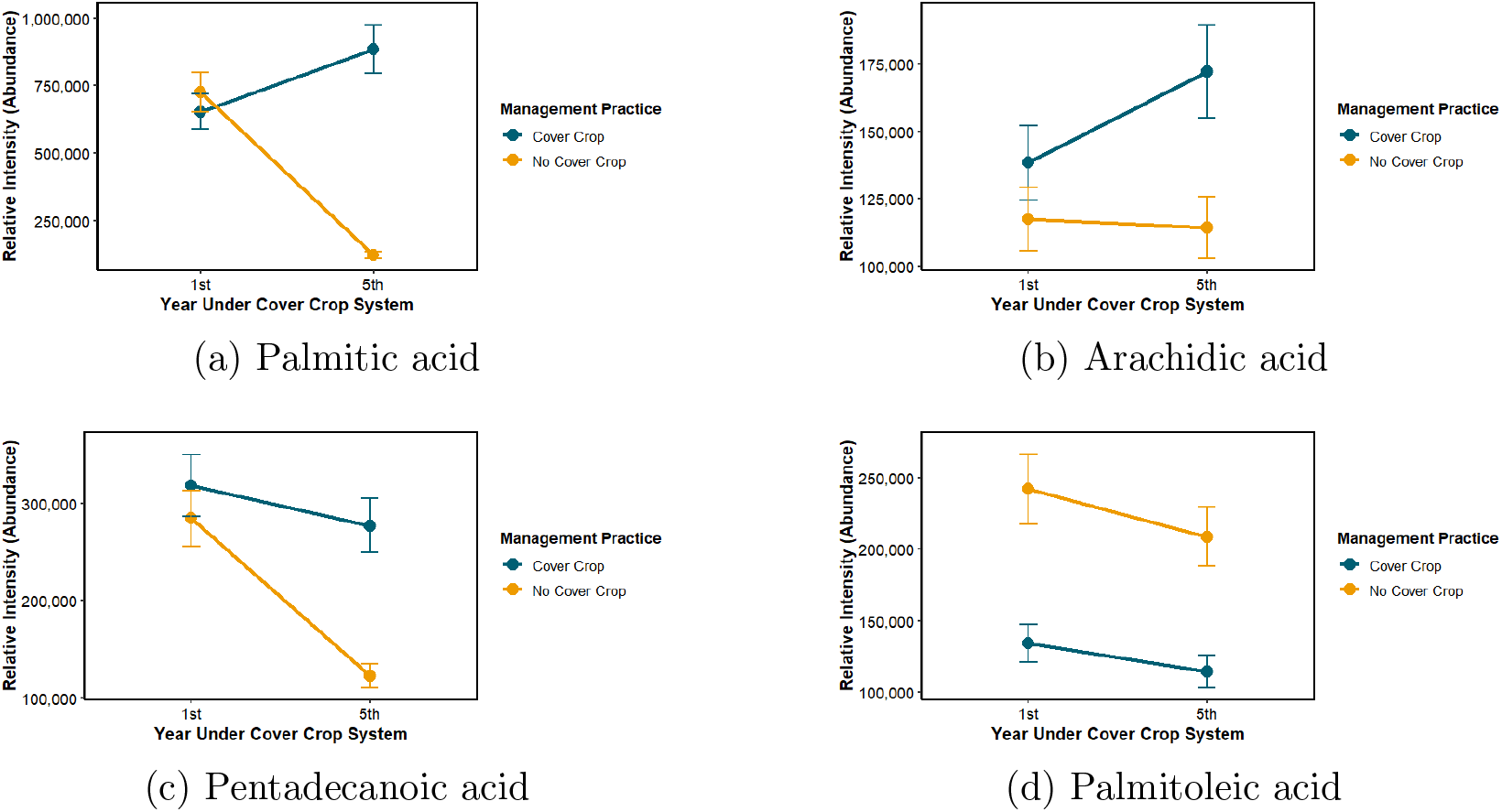
Relative intensity of select fatty acids. Palmitic acid (A) and arachidic acid (B) increased in the 5th-year cover crop (CC) treatment, while pentadecanoic acid (C) and palmitoleic acid (D) decreased over time in both treatments.

### 3.5. Metabolic Network Correlations

To explore the relationships and possible shared biochemical pathways among the most influential compounds, we constructed a Debiased Sparse Partial Correlation (DSPC) network (Figure 12). This analysis went beyond simple abundance and revealed a complex web of significant positive correlations among these key metabolites. The results indicated that the accumulation of these compounds was not random; instead, it is part of a coordinated metabolic response. Notably, several fatty acids, such as arachidic acid and eicosenoic acid, along with potent bioactive compounds like sclareol and phloretin, were highly interconnected. This suggests their production or degradation may be linked. In this complex network, Eicosapentaenoic acid emerged as a central hub, showing strong connections with many other metabolites. Its position suggests it may play a vital role in the soil’s metabolic response to long-term cover cropping. This could influence the abundance and activity of many other compounds, making it an important target for future mechanistic studies.

**Figure 12.**
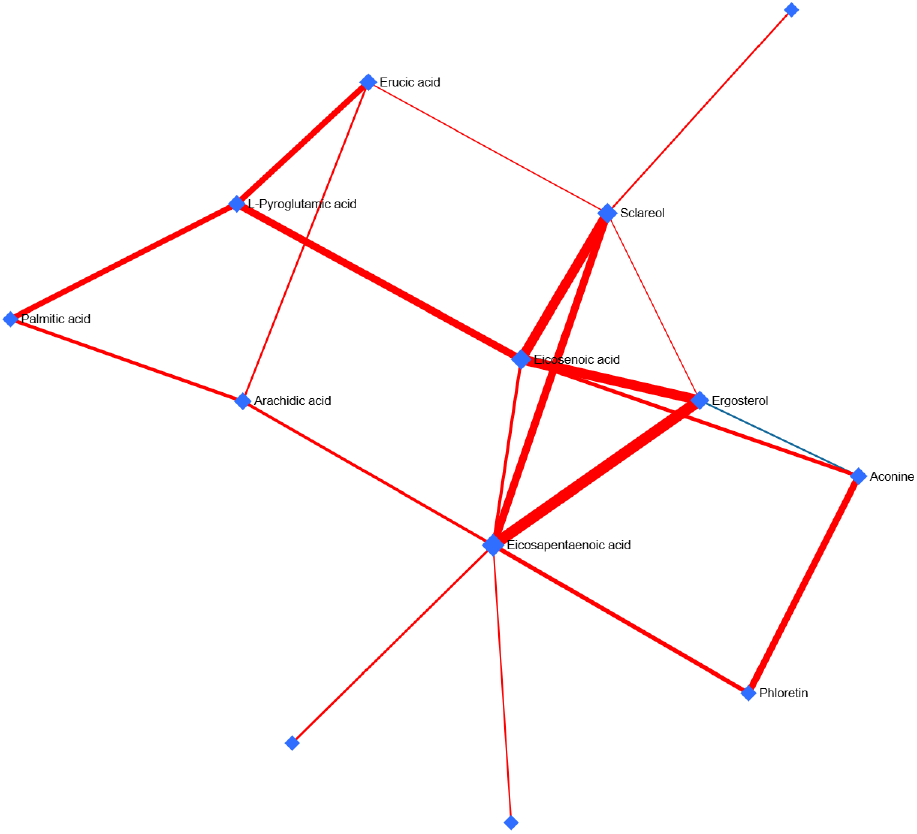
Debiased Sparse Partial Correlation (DSPC) network of key metabolites. Nodes represent metabolites, and edges represent significant positive correlations. The network highlights the interconnectedness of fatty acids and secondary metabolites, with eicosapentaenoic acid acting as a central hub.

## 4. Discussion

This study successfully used an untargeted metabolomics approach to reveal the complex chemical changes in the rhizosphere after long-term organic management with cover crops. Our main finding is that using cover crops significantly increases the diversity and abundance of bioactive metabolites in the soil compared to controls without cover crops. This rich and varied chemical profile offers a basis for understanding how cover cropping helps create disease-suppressive soils, which are essential for sustainable organic agriculture.

Metabolomic analysis revealed divergent trends among fatty acids that point to a dual impact of cover cropping on the soil microbiome. First, the accumulation of long-chain fatty acids like palmitic and arachidic acid in the 5th-year cover crop treatment is consistent with an increase in total microbial biomass (Frostegård et al., 2011). The increase in these compounds indicates that cover crop residues stimulate a larger and more active microbial community (Shu et al., 2022). This aligns with findings from a six-year study in an organic system where cover cropping frequency was the primary driver of increases in microbial biomass, even more so than compost additions (Brennan and Acosta-Martinez, 2017). This concept is central to “general suppression,” where a healthy microbial ecosystem competes with pathogens for resources and space, which reduces the occurrence of diseases caused by harmful organisms (Van Elsas et al., 2012). Second, the observed decrease in other fatty acids, such as pentadecanoic and palmitoleic acid, suggests a more complex restructuring of the microbial community over time, possibly indicating a shift in the dominant microbial phyla or a transition in community function as the soil ecosystem matures under continuous cover crop management.

Beyond fostering general suppression, our findings show that cover crops not only boost general microbial activity but also introduce key secondary metabolites that can help with “specific suppression” (Weller et al., 2002). The putative identification of compounds like the flavonoid hesperidin and the steroidal alkaloid tomatidine is especially important. Flavonoids are wellknown for their roles as signaling molecules in plant-microbe interactions and for their direct antimicrobial effects (Zawadzka et al., 2021). While hesperidin has moderate antimicrobial activity, its real effect may come from microbial biotransformation into its more active form, hesperetin. This process happens when plants face challenges from pathogens (Choi et al., 2022). Toma-tidine is a more complex example of a co-evolutionary arms race. It is the detoxification product of the strong antifungal *α*-tomatine. This conversion is mediated by the tomatinase enzyme from pathogens like *Fusarium oxysporum*, allowing them to overcome the plant’s chemical defense (Pareja-Jaime et al., 2008). However, this detoxification is not complete. While tomatidine is less toxic to the adapted pathogen, it still has significant antifungal activity against many other non-adapted fungi. In fact, it has been shown to be more effective than *α*-tomatine against yeast. This suggests that tomatidine helps create a broader suppressive environment (Ito et al., 2004, 2007). Additionally, the detection of the fungal sterol ergosterol, a reliable biomarker for living fungal biomass (Montgomery et al., 2000), indicates a dynamic fungal community. While compounds like sclareol and phloretin have also been recognized for their antimicrobial and allelopathic effects. These compounds contribute to the diverse chemical shield created by cover crops (Gallegos et al., 2016; Schlatter et al., 2017). The coordinated increase of these specific compounds, as highlighted in the metabolic network analysis (Figure 12), suggests the cultivation of a targeted chemical shield against pathogens, driven directly by the cover crop practice.

The results of this study are important for organic farming. By offering a detailed understanding of how cover crops work at the chemical level, farmers can go beyond a general method. They can start to choose cover crop mixtures that specifically address certain disease challenges. This contrasts with a focus on pesticides that aims for a single result (Couëdel et al., 2019). For example, a mixture of crucifers and legumes can provide biofumigation through glucosinolate compounds from the crucifer. At the same time, the legume can fix nitrogen and support beneficial arbuscular mycorrhizal fungi, stacking benefits and reducing the drawbacks of monoculture (Couëdel et al., 2019). Choosing species that produce certain allelochemicals or antifungal compounds could thus become a primary, non-polluting way to control diseases. This approach fits well with the principles of organic agriculture, which aim to reduce outside inputs and improve the natural resilience of the agroecosystem (Rigby and Cáceres, 2001).

While this study provides solid evidence for how changes in metabolites can help suppress diseases, there are some limitations to note. The compound identifications are based on two levels of confidence: high-confidence MS^2^ annotations based on fragmentation patterns and lower-confidence MS^1^ annotations based on accurate mass alone. While MS^1^ is a powerful tool for screening, these identifications are tentative and would need confirmation with authentic chemical standards to ensure accuracy. This study shows a strong connection between the enriched metabolic profile and the potential for disease suppression; however, future work should try to establish direct cause and effect. Future research should focus on carrying out bioassays that test these soil extracts against specific pathogens to link the observed chemical changes to pathogen inhibition. Also, isolating key identified compounds, like tomatidine and hesperidin, and testing their direct antimicrobial effectiveness in the lab would confirm their role in suppression. Combining soil metabolomics with metagenomic sequencing in future studies could directly connect changes in the soil’s chemical profile to specific shifts in the microbial community. This would help answer important questions about which organisms produce or are affected by these bioactive compounds.

## 5. Conclusion

This research demonstrates that long-term organic cover cropping cultivates a chemically rich and diverse soil environment. This altered metabolome, characterized by an increase in fatty acids linked to microbial biomass and the exclusive accumulation of potent antimicrobial compounds like tomatidine and sclareol, provides a clear chemical basis for the development of disease-suppressive soils. The findings provide molecular-level validation for the benefits of cover cropping, showing how this practice enhances both the broad, competitive capacity of the soil microbiome (“general suppression”) and improves the soil with a targeted chemical collection (“specific suppression”). These insights support a shift toward the strategic design of cover crop mixtures to achieve specific agronomic outcomes, offering a powerful, nature-based tool for sustainable agriculture.

## Acknowledgements

This work was supported by the USDA The Organic Agriculture Research and Extension Initiative (OREI) (Grant Number 2022-51300-37883), USDA/ARS Dale Bumpers Small Farm Research Center from the USDA Agricultural Research Service (Grant Number 58-6020-6-001), Center for Agroforestry at University of Missouri, and Lincoln University. We gratefully acknowledge Dr. Zhentian Lei and Dr. Lloyd Sumner at the University of Missouri Metabolomics Center for their assistance with the metabolomics analysis.

## Notes

### Competing Interest Statement

The authors have declared no competing interest.

### Summary of Updates

This version of the manuscript has been revised to update the coauthors and their affiliations.

